# Integrating carbon stocks and wildlife connectivity for nature-based climate solutions

**DOI:** 10.1101/2022.08.22.504302

**Authors:** Paul O’Brien, John S. Gunn, Alison Clark, Jenny Gleeson, Richard Pither, Jeff Bowman

## Abstract

Actions to protect against biodiversity loss and climate change will require a framework that addresses synergies between these interrelated issues. In this study we present methods for identifying areas important for the implementation of nature-based climate solutions and biodiversity conservation by intersecting high resolution spatial data for carbon storage and terrestrial connectivity. We explored the spatial congruence of carbon and connectivity in Ontario, Canada and examined effectiveness of current protected areas coverage. We found a weak positive relationship between carbon stocks and terrestrial connectivity; however, our maps revealed large hotspots, with high values of both indices, throughout the boreal forest and northern peatlands and smaller, isolated hotspots in the settled landscapes of the south. Location of hotspots varied depending on whether we considered forest or soil carbon. Further, our results show that current protected and conserved areas in Ontario only cover 13% of landscapes with the highest values for both carbon storage and connectivity. Protection or restoration of areas that maximize the co-benefits of carbon storage and connectivity would make significant contributions towards ambitious national targets to reduce greenhouse gas emissions and conserve biodiversity.

## Introduction

Trends in global climate change and biodiversity loss continue to be two of the most pressing threats facing humans and nature. Avoiding catastrophic outcomes requires immediate large scale, coordinated international, national, and subnational efforts. While strategies to address both global crises have traditionally been treated separately, interdependencies between the two suggest that these issues should be addressed holistically and synergistically. Nature-based solutions have emerged as a promising framework to meet both climate change and biodiversity goals.

Nature-based Solutions (NbS), as defined by the International Union for Conservation of Nature (IUCN), are actions to protect, manage, and restore natural ecosystems with the aims of addressing societal issues (e.g., mitigating climate change), while simultaneously providing benefits to human wellbeing and biodiversity (Cohen-Shacham, Walters, Janzen, & Maginnis, 2016). NbS is borne from the knowledge that sustainable ecosystem management, ecological restoration, and protected and conserved areas can reduce carbon emissions and increase carbon stored in plants and soil. The term NbS has been used as an umbrella for a range of actions including ecosystem-based adaptation, ecosystem restoration, natural climate solutions, green/blue infrastructure, and protected areas (Cohen-Shacham et al., 2019; Nesshöver et al., 2017; Seddon et al., 2020). Many government and non-government NbS efforts have focused on restorative actions, such as planting trees (Seddon et al., 2021). Ecosystem restoration is an important part of the solution; however, it should not be seen as a silver-bullet (Holl & Brancalion, 2020). While restoration efforts (e.g., reforestation of mixed, native forests) provide meaningful biodiversity benefits (Wang, Zhang, Li, & Wu, 2021) and increase carbon sequestration and storage (Lewis, Wheeler, Mitchard, & Koch, 2019), proactively conserving large intact ecosystems with high ecological integrity should remain a focus (Cook-Patton et al., 2021; Grantham et al., 2020; Locke et al., 2019; Noon et al., 2021). Protection of intact ecosystems can be more cost effective than restoration of degraded habitats (Cook-Patton et al., 2021; Drever et al., 2021; Watson et al., 2018) and provide multiple, synergistic benefits by maintaining existing carbon sinks and preventing large, potentially irrecoverable carbon emissions while providing biodiversity benefits (Arneth et al., 2020; Proctor, Schuster, Buxton, & Bennett, 2022). Protection can enable national and subnational strategies to more immediately maximize synergies between climate change mitigation and biodiversity goals, while also conserving other essential ecosystem services.

Canada is a signatory to both the United Nations Framework Convention on Climate Change (UNFCCC) and the Convention on Biological Diversity (CBD). In accordance with these international agreements, parties committed to keep global warming to within 1.5 – 2°C (Paris Agreement) and protect at least 17% of lands and 10% of inland waters by 2020 (CBD; Aichi Target 11). Canada has declared it will reduce greenhouse gas emissions by 40% by 2030 and reach net-zero emissions by 2050 through cuts in emissions, innovation and protection and restoration of lands and biodiversity (Emissions Reduction Plan; Government of Canada, 2022). Moreover, Canada has committed to protecting 25% of its land area by 2025 and 30% by 2030 (ECCC, 2020; CBD Post-2020 Framework; Secretariat of the CBD, 2020). Despite having some of the largest remaining intact forests (Watson et al., 2018; Wells, Dawson, Culver, Reid, & Morgan Siegers, 2020) and peatlands (Goldstein et al., 2020; Noon et al., 2021), both of which provide globally significant carbon storage and safeguarding for biodiversity (Harris et al., 2021; Kurz et al., 2013; Stralberg, Arseneault, et al., 2020; Wells et al., 2020), only 13.5% of terrestrial lands in Canada are currently protected (ECCC, 2021). Given Canada’s climate and biodiversity commitments and the urgency for action, there is great benefit to identifying areas that maximize co-benefits of climate change mitigation and biodiversity protection when planning for new protected and conserved areas as well as ecosystem restoration priorities.

One of the challenges of expanding protected and conserved areas coverage is determining which areas are most critical to protect (Carroll & Ray, 2021). With respect to climate change mitigation, this can be as straightforward as locating and subsequently protecting areas that store and sequester, or have the potential to sequester, a relatively large amount of carbon (Carroll & Ray, 2021; Goldstein et al., 2020). Identifying areas important for biodiversity is a more complicated undertaking given the many definitions and methods to measure biodiversity as well as the paucity of data in some regions and for particular species (Busch & Grantham, 2013; Soto-Navarro et al., 2020).

A commonly employed method to identify biodiversity hotspots has been to assess species richness or other species-based metrics (Marchese, 2015; Soto-Navarro et al., 2020). Key Biodiversity Areas (KBAs) is an emerging approach, guided by global and national standards, for identifying sites that are especially important for the conservation of biodiversity (Smith et al., 2019). However, protection of these biodiversity hotspots alone is not enough, and it is recognized that the design of well-connected networks of protected and conserved areas is integral to global and national biodiversity efforts. Landscape connectivity is critical for species movement and gene flow through dispersal and migratory movements (Noss et al., 2012) and reductions in connectivity have been found to be a strong driver of species extinctions (Hooftman, Edwards, & Bullock, 2016; Thompson, Rayfield, & Gonzalez, 2017). Maintaining or restoring landscape connectivity is cited as one of the most important biodiversity adaptation strategies, particularly in the face of climate change as species ranges shift to track suitable climates (Heller & Zavaleta, 2009; Schloss, Cameron, McRae, Theobald, & Jones, 2022).

Despite connectivity being essential for the long-term persistence of biodiversity (Ward et al., 2020) and recent advances in connectivity mapping (Dickson et al., 2019; Hall et al., 2021; Phillips, Clark, Baral, Koen, & Bowman, 2021), it is rarely incorporated into conservation planning (Carroll & Ray, 2021; Maxwell et al., 2020). Thus, we suggest that landscape connectivity should be better incorporated into planning for protected and conserved areas and priority restoration sites and integrated with areas important for carbon sequestration and storage. Together, carbon and connectivity attributes can provide a framework to consider synergies between climate change mitigation and biodiversity conservation.

Although there is an increasing recognition of the value of NbS in achieving climate change and biodiversity loss targets (IPBES, 2019; IPCC, 2022a, 2022b), few studies exist to aid national and regional efforts in identifying areas suitable for the implementation of nature-based solutions (but see Drever et al., 2021 and Zhu et al., 2021 for examples of national or regional NbS). Recent mapping efforts have identified hotspots for multiple ecosystem services across Canada, including carbon storage, freshwater, and recreational capacity (Mitchell et al., 2021). Although a KBA Canada Coalition is in the process of applying the global standards in Canada for terrestrial and freshwater areas, the results are not yet ready (http://www.kbacanada.org). Soto-Navarro et al. (2020) have produced maps highlighting global hotspots for biodiversity and carbon storage. Similarly, Dinerstein et al. (2020) identified globally important areas for protection of biodiversity and climate stabilization, while also incorporating an analysis of connectivity among existing protected areas. These studies are excellent for mapping broad-scale patterns of biodiversity and climate change stabilization. In Canada, however, the vast majority of public land is administered by the provinces, territories, and Indigenous governments (Carrol & Ray, 2021). Consequently, the creation of new protected and conserved areas and priority restoration sites requires coordination with provincial/territorial, Indigenous governments, local organizations, and regional land-use planners and will require more fine-scale resolution mapping resources.

Sothe et al. (2022) have produced a map of terrestrial carbon stocks (above and below-ground) in Canada at a resolution useful for regional, Indigenous, and local governments. These maps indicate that Canada possesses large stores of terrestrial carbon, much of which is considered irrecoverable and if lost would significantly contribute to atmospheric carbon levels (Noon et al, 2021). Recent advances have also been made with respect to connectivity mapping in Canada. Pither, O’Brien, Brennen, Hirsh-Pearson, and Bowman (2021) have used circuit theory (McRae & Beier, 2007) to produce the first pan-Canadian current density map, which reflects the probability of an animal moving through any point in a landscape and can be used to identify areas important for connectivity. This new connectivity map is also at a resolution useful for both national and regional land-use planners and has been validated using independent wildlife data. These recent advances in high-resolution mapping resources for carbon storage and landscape connectivity provide the opportunity for the first time to explore the spatial relationship between these two important values.

Here, we focus on Ontario, the second largest province in Canada, as a case study of how regional climate and biodiversity actions can support national targets. Ontario stands out as an area of interest because of the juxtaposition of large, intact natural areas and those significantly affected by human modification (Hirsh-Pearson, Johnson, Schuster, Wheate, & Venter, 2022) and because of its large carbon stocks (Sothe et al., 2022). Using high resolution spatial data, we employ methods to examine the intersection of existing carbon storage and connectivity maps to identify important areas for conservation, including those that may be priority for future restoration. We also explore the relationship of these two metrics by testing the hypothesis that since sites with high carbon stocks also tend to have low cost for animal movement, these high carbon sites should also contribute disproportionately to connectivity. We predicted a positive relationship between movement probability estimates from circuit theory and carbon stocks. We further predicted that areas most important for carbon storage and connectivity will be large intact landscapes.

## Methods

We used a bivariate mapping approach to explore the spatial congruence of carbon and connectivity layers in Ontario and examine how effective current protected areas coverage is at conserving hotspots between these layers. We then use the resulting maps to make recommendations for areas in Ontario that would be most effective at maximizing synergies between carbon storage and connectivity, and thus should be considered high priorities for protection.

### Carbon Data

As a measure of climate change mitigation value, we used a recently created 250-m resolution map of terrestrial carbon stocks of Canada (Sothe et al., 2022). The researchers produced separate maps of existing forest carbon and soil carbon stocks. The forest carbon layer includes estimates of above-ground live biomass, dead plant matter, and below-ground root biomass and the soil organic carbon layer provides estimates of carbon stocks for the top 1-m depth across Canada. Both the forest and soil carbon stock maps were produced using a combination of field measurements, satellite data, climate and topographic variables, and a machine learning algorithm. These maps represent the first seamless estimates of terrestrial carbon stocks in Canada at such a high resolution. For the present study we first cropped raster extents to that of Ontario and then resampled layers to a spatial resolution of 300 m to match that of the connectivity map. Due to the large differential in carbon stock sizes between the forest and soil carbon pools, we explored the relationship of each layer with connectivity separately. Sothe et al. (2022) reported that Canadian forests store approximately 18.4 Pg C, compared to the 306 Pg C of soil organic carbon (SOC) stored in the top 1 m. Given that carbon stocks contained within the soil carbon pool in Canada are vastly larger than that of the forest carbon pool, we decided to separate these two layers for the following analysis.

### Connectivity Layer

To identify areas important for connectivity, we used the 300-m resolution current density map of Canada produced by Pither et al. (2021). The authors modelled connectivity across Canada using a circuit theory approach, which draws on similarities between animal movement through a landscape and flow of electricity through a circuit, allowing for identification of multiple movement pathways. Electricity travels through a circuit according to a random walk, and this property allows animal movement as a random walk (Doyle & Snell, 1984) to be modeled using principles from electric circuits. Circuit theory models require a movement cost surface as an input, which represents anthropogenic and natural landscape features by the degree to which they impede or facilitate movement of animals. The cost surface used by Pither et al. (2021) was produced using the most up to date spatial data including the Canadian human footprint (Hirsh-Pearson et al., 2022) and a recently developed national road layer (Poley, Schuster, Smith, & Ray, 2022). In their study, the authors modelled terrestrial connectivity, such that natural features like forests and wetlands represented a low cost to movement, while roads and cities as well as large water bodies and mountains represented a high cost to movement (for more details see Pither et al., 2021). The resulting current density map output reflects the probability of movement of an animal through any given point across the landscape. Current density represents the amount of electrical current (measured in amps (A)) flowing through a given cell and is analogous to the probability of a random walker using that location. Current density in a 300-m pixel is proportional to the probability of that pixel being used during a random walk through the province. The authors used a wall-to-wall method of Circuitscape implemented in Julia (Hall et al., 2021, Phillips et al., 2021) to produce the first pan-Canadian, multi-species current density map at a resolution fine enough to support regional actions, such as for our provincial scale study.

### Comparing carbon and connectivity

To test the relationship between carbon storage and connectivity, we evaluated Spearman rank correlations between current density values and raster values of both forest and soil carbon. We used a bivariate mapping approach to identify important areas of overlap between current density and carbon layers. We calculated quantiles at 20% intervals for the current density, forest, and soil carbon layers and then reclassified cell values for each raster layer with a value of 1 - 5 based on which percentile range they fell within. We then overlapped the current density map with each carbon map to produce two bivariate maps for current density by forest carbon and current density by soil carbon. Cell values for the bivariate maps were calculated using all unique combinations of the two layers, which resulted in 25 different cell values (5 × 5 matrix; e.g., Soto-Navarro et al., 2020) varying in degree of importance for each layer. We followed a similar approach to Mitchell et al. (2021) and Soto-Navarro et al. (2020) such that areas where cells from both layers are within the top 20% represent areas most important for protection, while areas in the lowest 20% for both layers may indicate areas important for restoration. We use the phrase “hotspot” to refer to areas where the highest quantiles (top 20%) for both connectivity and carbon overlap.

### Overlap of existing protected areas with carbon and connectivity hotspots

To examine how existing protected areas overlap with hotspots for carbon storage and connectivity, we calculated the proportion of each quantile combination within protected areas boundaries. First, we calculated the area covered by each of the 25 quantile combinations. Then, using a vector layer of Ontario protected and conserved areas from the Canadian Protected and Conserved Areas Database (ECCC, 2021), we extracted raster values from each of the maps and calculated percent area protected within each group. Using anticipated targets for terrestrial land conservation based on draft content for the Post-2020 Global Biodiversity Framework (Secretariat of the CBD, 2020), we identified protection of 30% to be a goal for protection.

## Results

Scatterplots of current density values against both forest and soil carbon values in Ontario displayed weak positive relationships between connectivity and carbon storage (Fig. 1 and 2; Spearman rank correlation of forest carbon and current density: rho= 0.074, p < 0.001; correlation of soil carbon and current density: rho = 0.093, p < 0.001). Our bivariate maps revealed varying amounts of spatial overlap between current density and carbon in Ontario, depending on the location and the type of carbon stock. The spatial distribution of hotspots between current density and carbon varied for forest carbon (Fig. 3) and soil organic carbon (Fig. 4). There was low overlap of highest quantile areas between connectivity and forest carbon (i.e., 4.6% of the map consisted of overlapping highest quantile areas, or hotspots) and only 3.4% between connectivity and soil carbon. Further, there was only a 2% overlap between hotspots identified by the connectivity-forest carbon map (Fig. 3) and the connectivity-soil carbon map (Fig. 4; Fig. S1).

**Fig 1.**
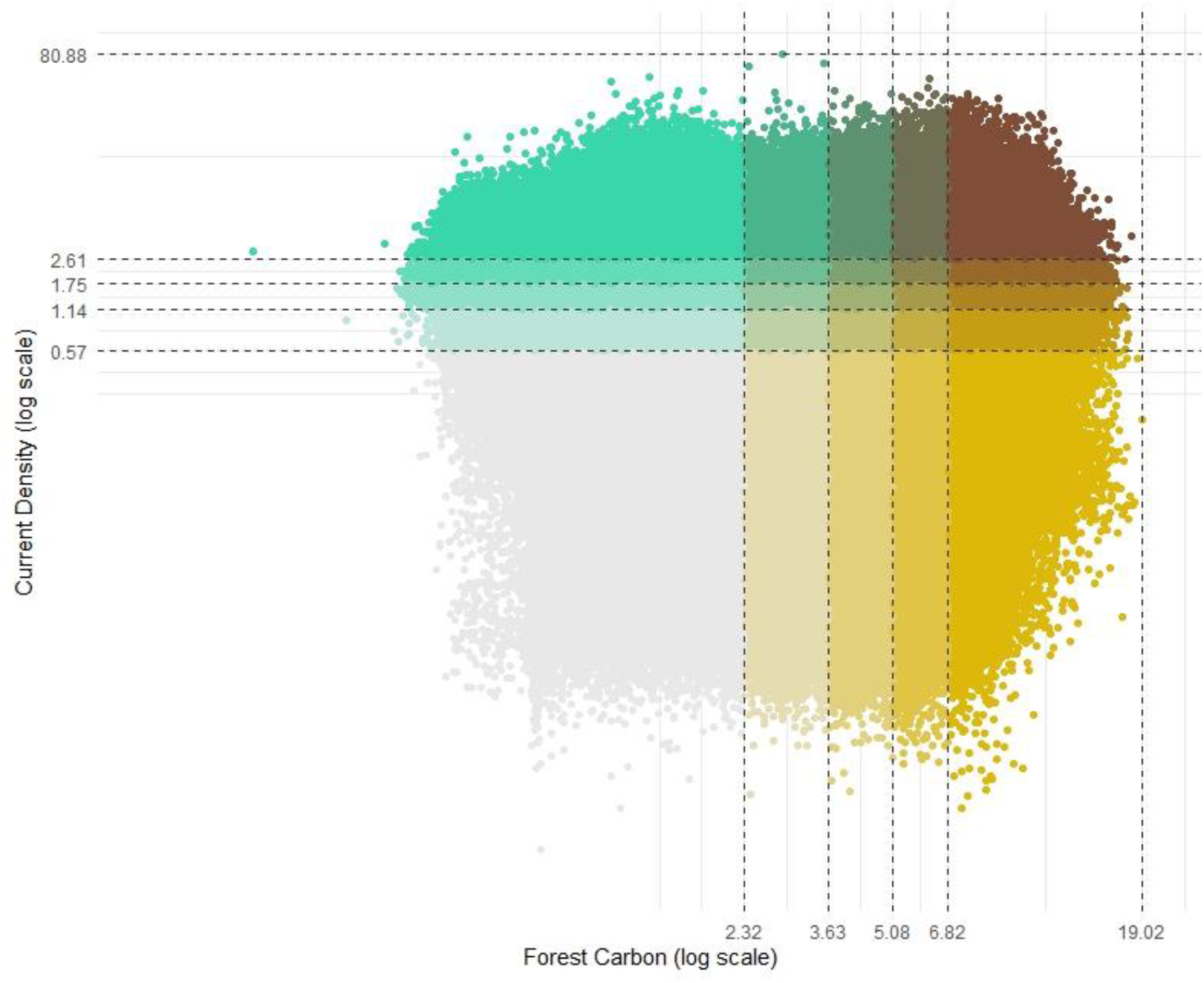
Scatterplot of cell values from forest carbon and current density, representing the probability of animal movement, raster layers for Ontario. Values are divided into quantile groups (by 20% intervals) with the colour gradient expressing overlap between percentile groups for both datasets. Points in the top right corner (dark brown) represent areas where cell values for both layers are in the top 20% quantile representing high connectivity and high carbon storage.

**Fig. 2.**
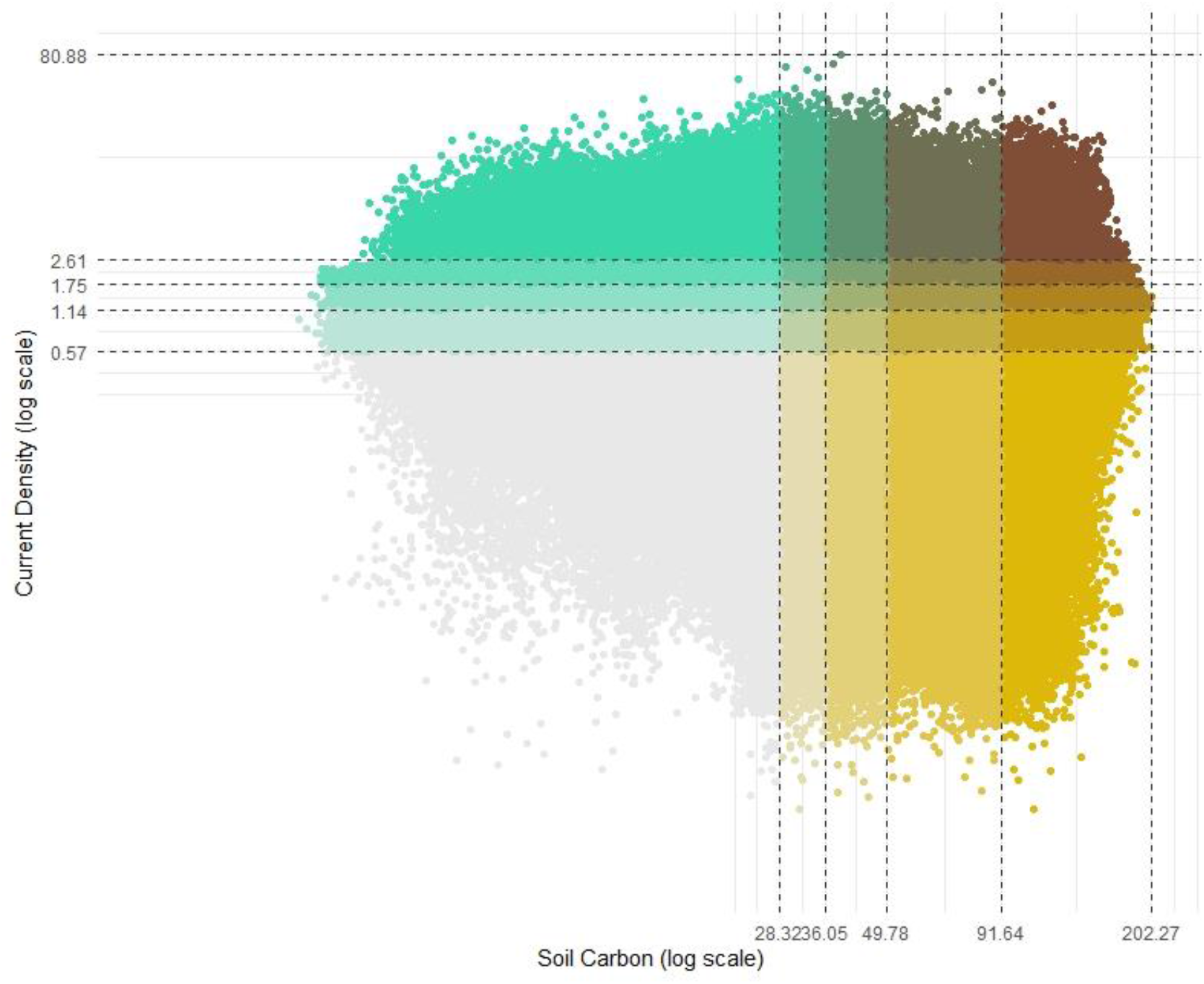
Scatterplot of cell values from soil carbon and current density, representing the probability of animal movement, raster layers for Ontario. Values are divided into quantile groups (by 20% intervals) with the colour gradient expressing overlap between percentile groups for both datasets. Points in the top right corner (dark brown) represent areas where cell values for both layers are in the top 20% quantile representing high connectivity and high carbon storage.

**Fig. 3.**
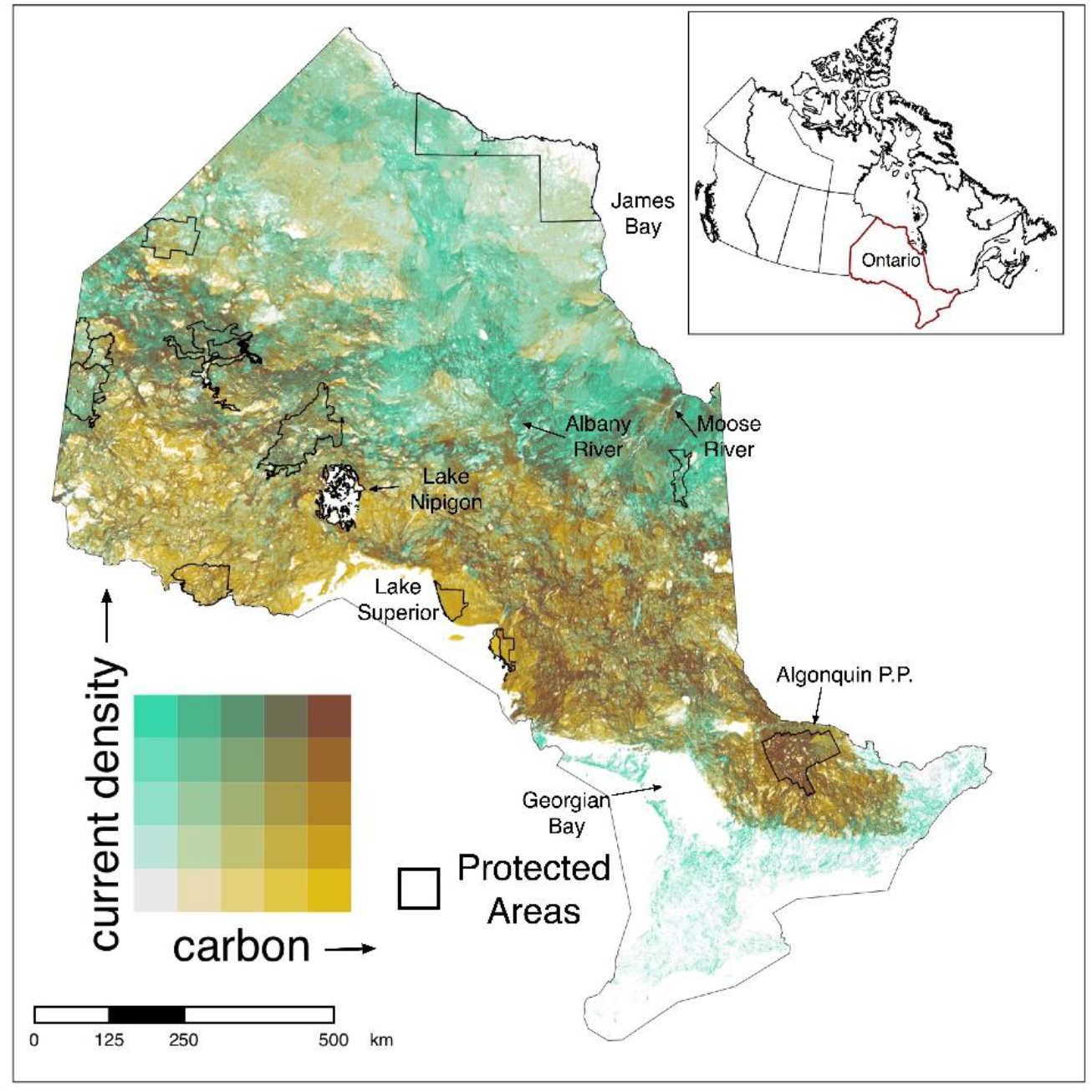
Bivariate map showing spatial overlap between current density, representing the probability of terrestrial animal movement, and forest carbon biomass in 300m cells across Ontario. Colour scales are based on quantile intervals (in 20% increments). Dark brown regions (top right corner of colour matrix) represent cells that are in the top 20% quantile for both current density and carbon layers. Solid black lines show the boundaries of selected large (> 150000 ha) protected areas (excluding marine protected and conserved areas).

**Fig. 4.**
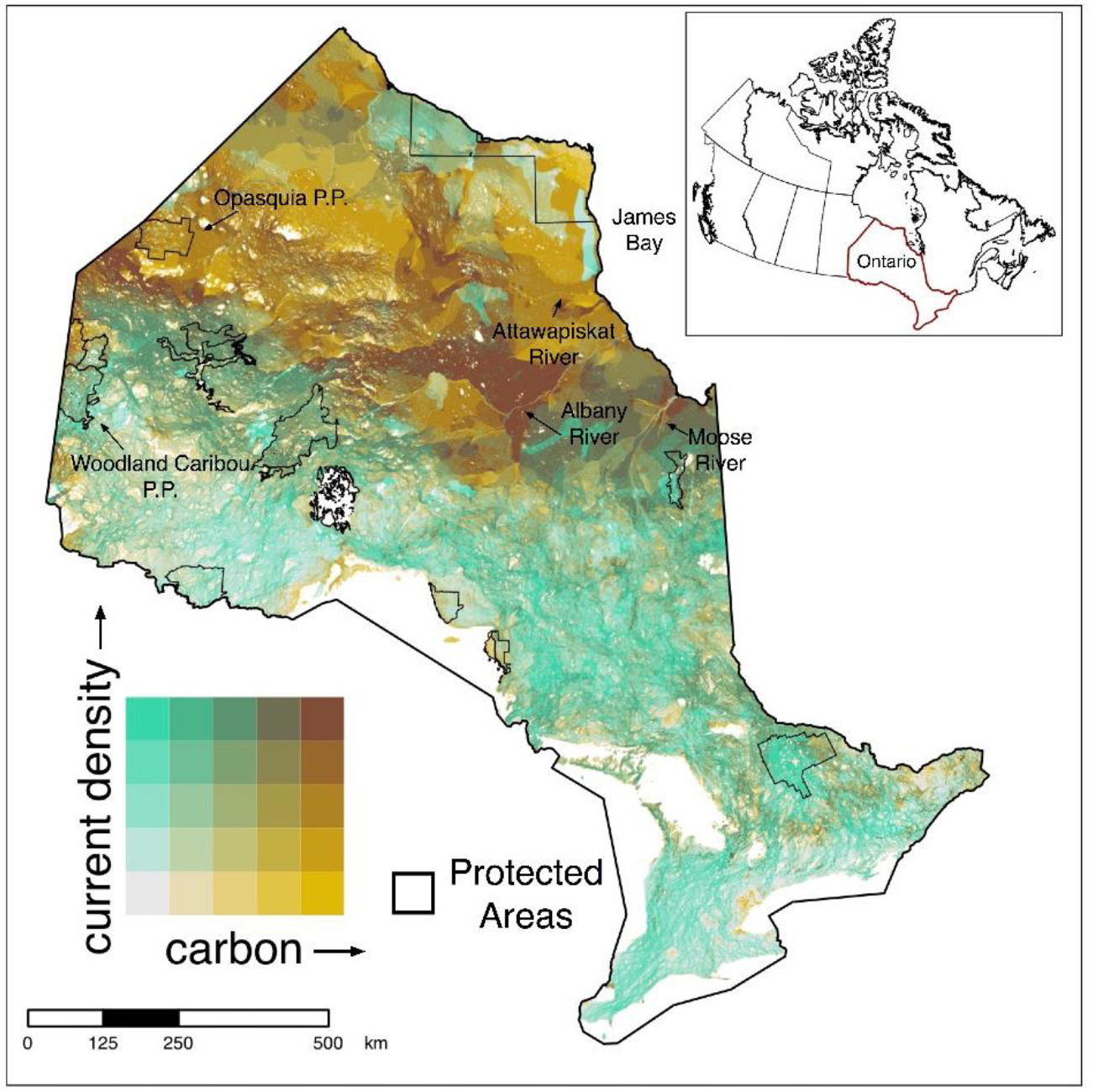
Bivariate map showing spatial overlap between current density, representing the probability of animal movement, and soil organic carbon biomass in 300 m cells across Ontario. Colour scales are based on quantile intervals (in 20% increments). Dark brown regions (top right corner of colour matrix) represent cells that are in the top 20% quantile for both current density and carbon layers. Solid black lines show the boundaries of selected large (> 150000 ha) protected areas (excluding marine protected and conserved areas).

Areas most important for both connectivity and forest carbon storage were largely found within the northern regions of the province, which coincides with the Boreal Shield ecozone of Canada. For example, prominent areas of overlap occur around Algonquin Provincial Park, north of Lake Superior, and moving northwest from Lake Nipigon into eastern Manitoba (Fig. 3). Fewer hotspots were evident farther north in Ontario; however, important areas did appear along rivers south of James and Hudson Bay, such as the Moose and Albany rivers (Fig. 3, Fig. 5a). In the south, hotspots for connectivity and forest carbon were concentrated in the area east of Algonquin Park towards the Canada-US border, which is part of the existing Algonquin to Adirondacks (A2A) connectivity initiative (Fig. 5b). Our analysis did not identify any hotspots in the southern region of the province; however, important corridors still exist in the southwest despite the high degree of anthropogenic disturbance. For example, areas of high current density can be seen along a north-south corridor east of Georgian Bay (Fig. 5b). These areas of high connectivity and low carbon storage may be important to protect to maintain connectivity and provide good opportunities for restoration (e.g., tree planting) to enhance both connectivity and carbon stocks.

**Fig. 5.**
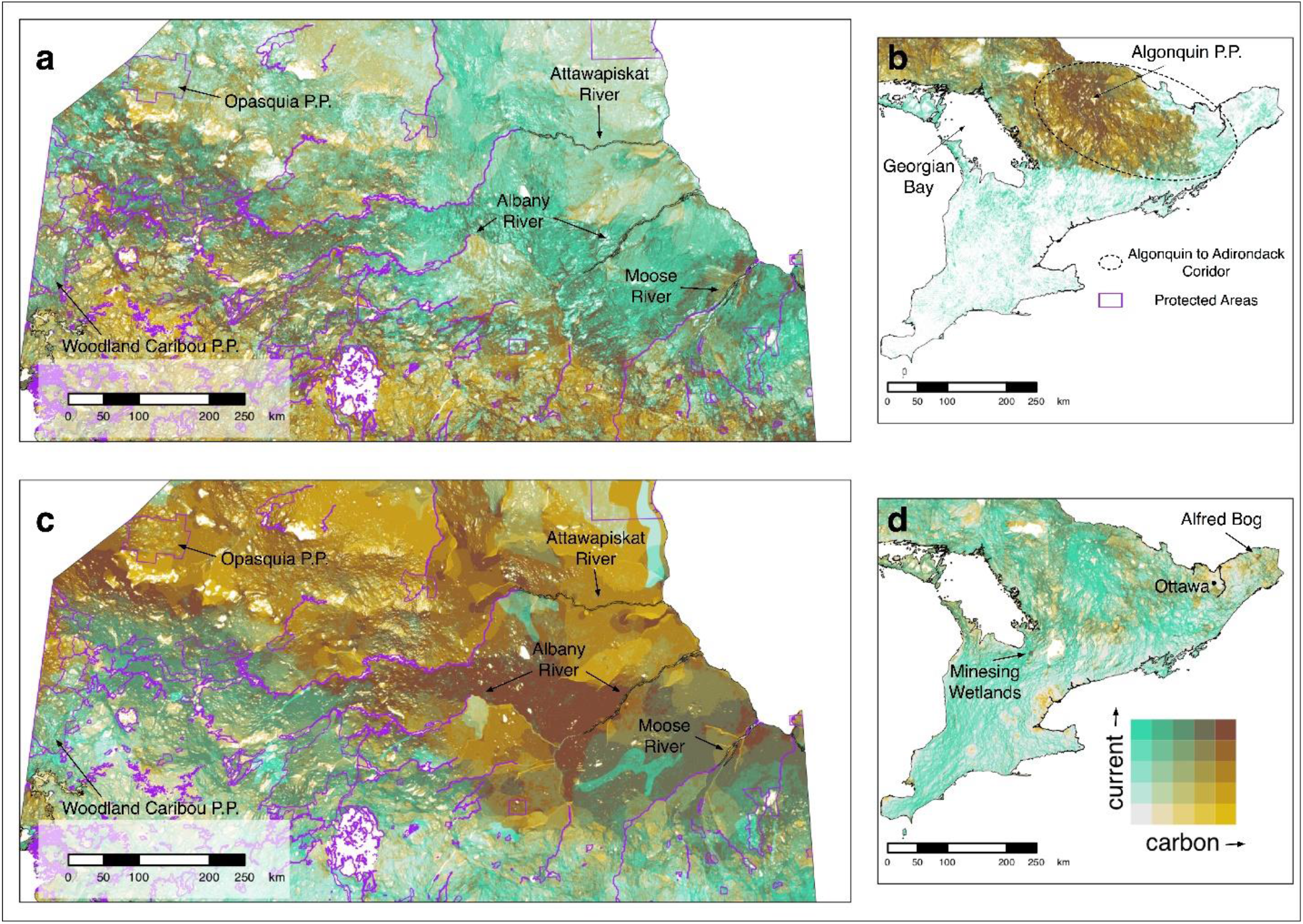
Vignettes to highlight hotspots for current density, representing the probability of animal movement and: (a) forest carbon in northern Ontario, (b) forest carbon in southern Ontario, (c) soil carbon in northern Ontario, and (d) soil carbon in southern Ontario. Purple lines show the boundaries of protected and conserved areas in Ontario and the dashed oval represents the general extent of the Algonquin to Adirondacks corridor. Protected areas boundaries were excluded from panels (b) and (d) to maintain map clarity.

Hotspots for connectivity and soil carbon storage covered a large area across Ontario (∼3.7 million hectares total) with many of these areas occurring throughout the Hudson Bay Lowlands in northern Ontario. The most important areas were found south of James Bay around the mouth of the Moose River, between the Albany and Attawapiskat Rivers, and nearing the Ontario-Manitoba border between Woodland Caribou and Opasquia Provincial Parks (Fig. 4, Fig. 5c). These areas are particularly important as they are also identified as important areas for connectivity and forest carbon. Many smaller hotspots (top 20% values for both layers) were also identified throughout the north as well as large areas where top 20% values for one layer overlap with top 40% values of another. Similarly, several smaller areas appear in southern Ontario where top 20% values for current density overlap with top 40% values for soil carbon (e.g., Minesing wetlands and Alfred Bog; Fig. 5d).

Our analysis indicates that the current system of protected areas in Ontario protects 12% of hotspots for connectivity and forest carbon and only 1% of hotspots for connectivity and soil carbon (Fig. 6). The highest percentage of area protected for overlap of connectivity and both forest and soil carbon were found in areas most important for carbon, but least important for connectivity, at 19.7 and 22.2 %, respectively (Fig. 6).

**Fig. 6.**
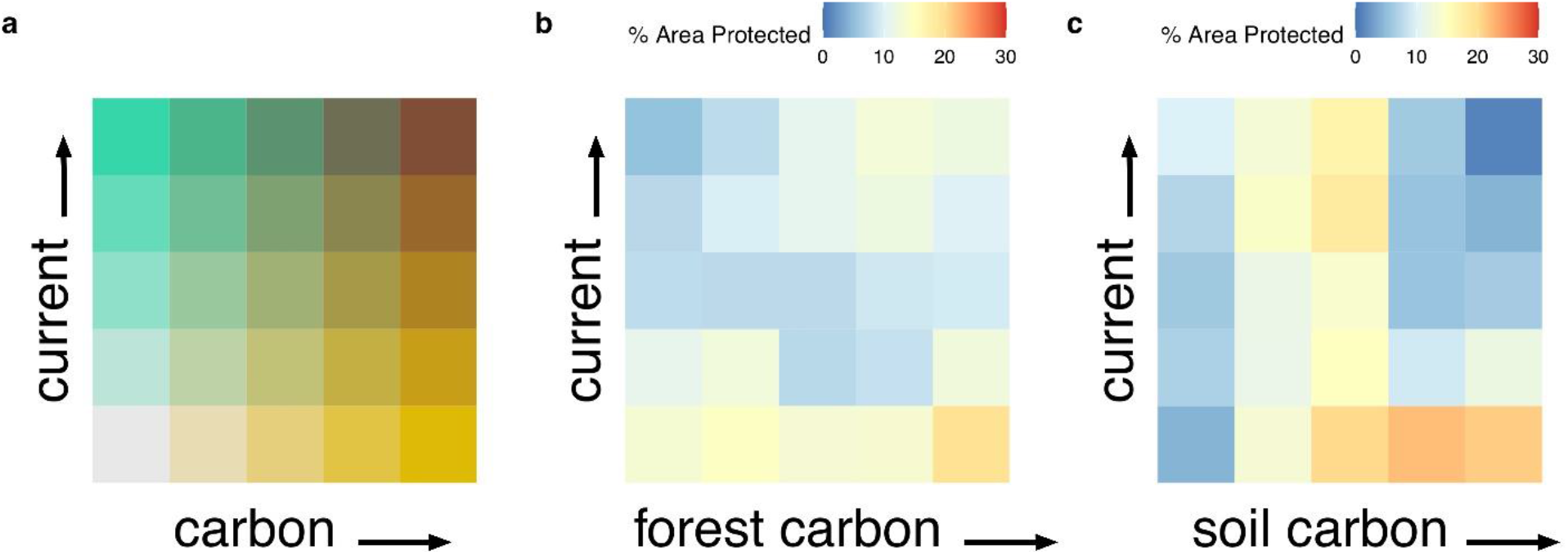
Colour matrices displaying (a) the underlying layout for the spatial overlap of current density and carbon layers with cell values divided in 20% quantile intervals; the percent area within each quantile group contained within current protected areas for bivariate maps of current density and (b) forest carbon and (c) soil organic carbon.

## Discussion

We found a low degree of geographic overlap between the highest quantile areas for connectivity and forest carbon (4.6%) and between connectivity and soil carbon (3.4%). We also found overall connectivity to be weakly correlated with both forest and soil carbon storage. This is consistent with other studies reporting a weak correlation between carbon and biodiversity (Di Marco, Watson, Currie, Possingham, & Venter, 2018) and climate refugia (Carroll & Ray, 2021). Although pixels with high carbon storage values were often areas with low movement costs, this did not always translate into high current densities, largely due to the influence of geometry and the tendency for proximate high-cost areas to lead to a funneling of current flow (Marrotte et al., 2017).

The limited spatial alignment of the highest quantiles for carbon and connectivity suggest that the hotspots where the top 20% for both measures overlap may be priorities for protection.

Further, we found there was only a 2% overlap between hotspots identified between the two maps (i.e., hotspots for connectivity, forest carbon, and soil carbon). These overlapping areas between the two maps (Figs. 3 and 4) were all found in the naturally intact landscapes of northern Ontario: south of Opasquia Provincial Park at the Ontario-Manitoba border, along the Albany River, and at the mouth of the Moose River (Fig. S1). The limited overlap between these three metrics indicates that these areas are high priority for protection and that conservation strategies with an objective to incorporate both connectivity and carbon should consider forest carbon and soil carbon separately. Including only one or the other likely would result in areas of importance being missed, while pooling both together would lead to a bias towards regions important for soil carbon storage driven by the greater magnitude of carbon stored in soils (Crowther et al., 2019).

Ontario’s soil carbon stocks are largely contained within the vast boreal peatlands of the Hudson Bay Lowlands (Harris et al., 2021; Sothe et al., 2022) which is also where our bivariate maps reveal strong spatial congruence between landscape connectivity and soil carbon storage. These carbon stores have been identified as irrecoverable at a global scale meaning they would not be recoverable within the timeframe necessary to reach net-zero (Goldstein et al., 2020; Noon et al., 2021; Packalen, Finkelstein, & McLaughlin, 2014). The largest areas of overlap occurred in the northern part of the province where boreal forest and peatland ecosystems remain largely intact. Protection of these hotspot areas would make a considerable contribution towards climate change targets while enhancing biodiversity. Further, across northern Canada, especially northern Ontario, there is a high degree of overlap between Indigenous communities and territories and important carbon sinks (Townsend, Moola, & Craig, 2020); most of the hotspots identified by our analysis follow this trend. Thus, identification and design of protected areas in these regions should occur in collaboration with Indigenous communities/governments and to be guided by Indigenous expertise. Importantly, mapping resources such as we present here could be helpful for Indigenous governments to advance Indigenous Protected and Conserved Areas (IPCAs) efforts (Indigenous Circle of Experts, 2018).

Hotspots of soil carbon and connectivity appear in the region between the Albany and Attawapiskat rivers where a large east-west corridor coincides with these large, intact peatlands. Not surprisingly, this corridor has been shown to be critical to migrating species such as woodland caribou (OMECP, 2020; OMNRF, 2019) and birds (ECCC, 2013) and this area may provide a natural buffer or refugia for the persistence of biodiversity as the climate changes (Morelli et al., 2020; Stralberg, Arseneault, et al., 2020; Stralberg, Carroll, & Nielsen, 2020). Our maps also identify hotspots for both forest and soil carbon with connectivity at the western border of Ontario and Manitoba. Protection here would not only maximize carbon storage benefits (i.e., protecting important forest and soil stocks), but would also contribute to global biodiversity goals of creating well-connected networks of protected areas by connecting Woodland Caribou Provincial Park to the south with Opasquia Provincial Park to the north.

Isolation of Ontario’s north has safeguarded these hotspots to date; however, soil and peat stocks are increasingly threatened by land-use and land cover changes, such as mineral extraction, draining for agriculture, peat extraction, and shrubification (Harris et al., 2021). Due to the likely irrecoverability of soil carbon in this region, its importance to wildlife, and the potential to support biodiversity in the future warmer climate, protection of these hotspots could be critically important while also being cost-effective (Cooke-Patton et al., 2021; Goldstein et al., 2020). Currently only ∼1% of these hotspots are protected. Increasing protection in this region to even 5% has the potential to conserve ∼0.2 Pg of carbon, which is equivalent to CO_2_ emissions from 188 coal-fired power plants for a year (USEPA, 2015), while also helping to maintain connectivity between Canada’s eastern and western provinces (Murray et al., 2017). Given the high degree of overlap between Indigenous territories and carbon sinks (Townsend et al., 2020) and the importance of Indigenous managed-lands as habitat for biodiversity (Schuster, Germain, Bennett, Reo, & Arcese, 2019), these northern hotspots may be particularly important candidates for IPCAs. These should be implemented with consent and in partnership with Indigenous peoples and governments and should help advance Indigenous-led conservation.

Forest carbon makes up a smaller proportion of carbon stocks in Ontario. Forest carbon is generally considered recoverable carbon due to natural regeneration, reforestation activities, and the ability to store carbon long-term in harvested wood products. Our bivariate maps reveal areas with strong spatial congruence between landscape connectivity and forest carbon storage throughout the boreal shield ecozone of Ontario. Identified hotspots for connectivity and forest carbon may face more immediate threats from resource extraction (e.g., logging and mining), forest fires, pest outbreaks, extreme drought, and invasive species (Wells et al., 2020), with the latter four likely to increase as a consequence of climate change (Allen, Breshears, & McDowell, 2015; Anderegg et al., 2020; Price et al., 2013). Ontario has some of the largest intact boreal forests in Canada (Watson et al., 2018), but deforestation has occurred both within forestry tenures (Smith & Cheng, 2016; OMNRF, 2021) as well as outside of managed forests. While there are threats facing forest carbon stocks, the significance of soil carbon stocks in the north compared to forest stocks clearly expresses the disproportionately important role of soil carbon in climate change mitigation and the critical need to protect these stocks. Protection and management of these different carbon pools may therefore require different strategies.

The exceptional value of intact forests for carbon storage and biodiversity is generally noted meaning that protection of these areas is important (Watson et al., 2018). The recoverable nature of forests and their broad utility for connectivity could be more compatible with a combination of protection and improved forest management practices (Malcolm, Holtsmark, & Piascik, 2020). For example, an analysis of natural climate solutions in Canada found improved forest management (e.g., a combination of old growth conservation and regeneration after harvest) to be the fourth largest opportunity for climate change mitigation potential in 2030 (Drever et al., 2021). Aside from national and provincial protected and conserved areas, those areas that are managed in ways that support biodiversity and climate even when not the primary objective may be particularly effective at conserving and increasing connectivity in managed landscapes (e.g., forests; Maxwell et al., 2020).

In the southern part of the province, we identified spatial congruence between forest carbon and connectivity coinciding with the known Algonquin to Adirondacks Corridor (A2A). Addition of protected and conserved areas within this corridor could provide significant benefits to the transboundary movement of wildlife between Algonquin Park in Ontario, Canada, and Adirondack Park in New York State, U.S.A. Additionally, Proctor et al. (2022) found that protection of species at risk habitat in Ontario may be most cost effective in the central region of the province where land cost is low, but species at risk richness is still relatively high. This aligns with areas important for connectivity and forest carbon storage identified by our analysis.

Moreover, the eastern section of A2A and regions of southwestern Ontario exhibit importance for connectivity but are lacking in forest carbon stocks. These regions could therefore be an important focus of ecological restoration efforts (e.g., tree-planting initiatives, grassland restoration). Restoring degraded ecosystems and corridors is an imperative step to enhancing biodiversity and can increase both above- and below-ground carbon stocks (Gunn, Ducey, & Belair, 2019; Soto-Navarro et al., 2020; Valach et al., 2021). In this case, restoring within the A2A could help maintain connectivity, while also building carbon stores. Our maps could also be utilized to identify other areas where carbon stocks are high and restoration activities could reduce their vulnerability or likewise, where carbon is low and restoration could increase these values over time.

Our maps identified Catchacoma Forest, an old growth eastern hemlock (*Tsuga canadensis*) forest northeast of Lake Simcoe, as an area of interest for connectivity and forest carbon. In addition, Catchacoma Forest has been identified as the largest known old growth eastern hemlock stand in Canada and an important site for various species at risk (Quinby, 2020); however, it is threatened by the invasive hemlock woolly adelgid (*Adelges tsugae*; NRCAN, 2015) and is an actively logged area. We also identified small hotspots for connectivity and soil carbon in the south. Of particular interest are Minesing Wetlands and Alfred Bog, which represent some of the last large wetlands in southern Ontario. The Minesing Wetlands are currently being assessed as a Key Biodiversity Area, which further supports it as a priority area for protection. Similarly, Alfred Bog is a provincially significant wetland, an Area of Natural and Scientific Interest, has protection through various NGOs, and is in the process of Provincial Park designation (NCC, 2021). Protection of these provincially significant wetlands would provide multiple benefits for climate change mitigation, biodiversity, human wellbeing, as well as other important ecosystem services.

Interestingly, the results of our provincial analysis exhibit similarities with recent global analyses. Soto-Navarro et al. (2020) assessed the spatial overlap of global carbon stores with measures of reactive (vulnerability) and proactive (intactness) biodiversity. Similarly, Dinerstein et al. (2020) mapped global conservation priorities for biodiversity and climate stabilization as well as identified potential wildlife corridors between current protected areas. All three studies identify priority areas within the Hudson Bay Lowlands and Boreal Shield in Ontario, with few areas of importance occurring in southern Ontario. The high resolution of our analyses, however, allowed us to distinguish more fine-scale patterns of spatial overlap between carbon storage and connectivity, which will be useful for land-use planners across different jurisdictions. Moreover, the separation of forest and soil carbon in our study illustrated the differences in conservation priorities identified when assessing the relationship of these different carbon pools with landscape connectivity. Still, it is apparent from the current and previous studies that the Hudson Bay Lowlands is a hotspot for carbon storage, connectivity, and biodiversity (Dinerstein et al., 2020; Soto-Navarro et al., 2020) and is therefore a high priority for protection.

Existing protected and conserved areas in Ontario cover 10.7% of the province’s area (Ontario Parks, 2021). Our analysis indicates that 12% of hotspots (top 20% overlap) for connectivity and forest carbon and only 1% of hotspots for connectivity and soil carbon are contained within current protected areas. A lack of protection was also evident for hotspots of biodiversity and carbon globally (Soto-Navarro et al., 2020) and in Asia (Zhu et al., 2021). Similarly, Proctor et al (2022) found that 50% of species at risk in Ontario had less than 10% of their habitat protected by existing protected areas. A recent temporal analysis by Maxwell et al. (2020) revealed that expansion of protected areas (between 2010 and 2019) made minimal contributions to conserving various elements of biodiversity and ecosystem services, including carbon storage and connectivity. These findings stress the critical need for resources such as the above-mentioned studies and our current study for helping to guide the expansion of effective protected and conserved areas. Our integration of spatial data on carbon storage and connectivity provides a framework to consider synergies between climate change mitigation and biodiversity conservation, such that we were able to identify key areas for targeted NbS strategies in Ontario.

We believe our maps will be useful for land-use planners and stakeholders in identifying areas important for expanding protected areas; however, we acknowledge several caveats. We used the most up to date carbon maps available but there remain uncertainties associated with these carbon stock estimates, particularly in peatland soils (Sothe et al., 2022). Second, there may be uncertainty with how well the connectivity map we used predicts areas important for connectivity in northern Ontario given that much of the mammal data used to validate the map was from western Canada. Finally, while we would have ideally included data layers for carbon, connectivity, and biodiversity, the KBA data for Ontario was not yet available, however our maps still identified areas that coincided with designated and potential KBAs. Therefore, we suggest that future analyses could update our maps with biodiversity data, especially KBA locations as they become available. Despite these caveats, we consider that our maps accurately identify hotspots for connectivity and carbon storage in Ontario and can be a valuable resource for the identification of new protected and conserved areas and sites for ecological restoration.

## Conclusion

Global biodiversity and climate targets will require ambitious national and regional actions to protect and sustainably manage remaining areas of high ecological integrity and restore damaged and degraded areas to improve biodiversity. We intersected high-resolution spatial layers for carbon storage and terrestrial connectivity to provide a framework for exploring synergies between climate change mitigation and biodiversity conservation in Ontario, Canada. We found a weak relationship between these metrics; however, the maps we produced still revealed hotspots with high values for both connectivity and carbon storage in the province. Our results align with previous global analyses, providing further support that the areas identified are priorities for protection. We believe our high-resolution mapping resources will be useful for land-use planners and policymakers to identify areas important for nature-based climate solutions. Further, our methods could easily be adopted by other provinces, territories, and other jurisdictions to contribute to recent ambitious Canadian national targets to reach 30% protection of terrestrial lands by 2030. Critical to achieving the ambitious 30x30 target is not only increasing coverage, but also effectiveness of protected areas. Our approach can help to maximize the co-benefits for climate change mitigation and biodiversity conservation.

## Author Contributions

J.B., A.C., and J.G. conceived the study. J.B., J.S.G., and P.O. developed the methodology. P.O. was the lead for spatial analyses, mapping, and figures. All authors contributed to writing, reviewing, and editing of the manuscript.

## Acknowledgements

Financial support for this study was provided by the Ontario Ministry of Natural Resources, and Forestry for P.O., J.B., J.G., and A.C., New Hampshire Agricultural Experiment Station for J.S.G., and Environment and Climate Change Canada for R.P.

## Data Availability

Current Density Map and Movement Cost Layer (https://osf.io/z2qs3/; DOI 10.17605/OSF.IO/Z2QS3)

Forest Carbon Stock Map (https://doi.org/10.4121/14572929.v1) and Soil Carbon Stock Map (https://doi.org/10.4121/16686154.v3)

Our output Forest Carbon-Connectivity and Soil Carbon-Connectivity Maps (Dryad repository doi: XXXX)

## Literature Cited

Allen, C.D., Breshears, D.D., & McDowell, N.G. (2015). On underestimation of global vulnerability to tree mortality and forest die-off from hotter drought in the Anthropocene. Ecosphere 6(8):1–55.

Anderegg, W.R.L., Trugman, A.T., Badgley, G., Anderson, C.M., Bartuska, A., Ciais, P., … Randerson, J.T. (2020). Climate-driven risks to the climate mitigation potential of forests. Science 368(6497).

Arneth, A., Shin, Y.J., Leadley, P., Rondinini, C., Bukvareva, E., Kolb, M., … Saito, O. (2020). Post-2020 biodiversity targets need to embrace climate change. Proceedings of the National Academy of Sciences of the United States of America 117(49):30882–30891.

Busch, J., & Grantham, H.S. (2013). Parks versus payments: reconciling divergent policy responses to biodiversity loss and climate change from tropical deforestation. Environmental Research Letters 8(3):034028.

Carroll, C., & Ray, J.C. (2021). Maximizing the effectiveness of national commitments to protected area expansion for conserving biodiversity and ecosystem carbon under climate change. Global Change Biology 27(15):3395–3414.

Cohen-Shacham, E., Andrade, A., Dalton, J., Dudley, N., Jones, M., Kumar, C., … Walters, G. (2019). Core principles for successfully implementing and upscaling Nature-based Solutions. Environmental Science and Policy 98:20–29.

Cohen-Shacham, E., Walters, G., Janzen, C., & Maginnis, S. (2016). Nature-based solutions to address global societal challenges. Gland, Switzerland.

Cook-Patton, S.C., Drever, C.R., Griscom, B.W., Hamrick, K., Hardman, H., Kroeger, T., … Ellis, P.W. (2021). Protect, manage and then restore lands for climate mitigation. Nature Climate Change 11(12):1027–1034.

Crowther, T.W., van den Hoogen, J., Wan, J., Mayes, M.A., Keiser, A.D., Mo, L., … Maynard, D.S. (2019). The global soil community and its influence on biogeochemistry. Science 365(6455).

Di Marco, M., Watson, J.E.M., Currie, D.J., Possingham, H.P., & Venter, O. (2018). The extent and predictability of the biodiversity–carbon correlation. Ecology Letters 21(3): 365–375.

Dickson, B.G., Albano, C.M., Anantharaman, R., Beier, P., Fargione, J., Graves, T.A., … Theobald, D.M. (2019). Circuit-theory applications to connectivity science and conservation. Conservation Biology 33(2):239–249.

Dinerstein, E., Joshi, A.R., Vynne, C., Lee, A.T.L., Pharand-Deschênes, F., França, M., … Olson, D. (2020). A “global safety net” to reverse biodiversity loss and stabilize earth’s climate. Science Advances 6(36).

Doyle, P.G., & Snell, J.L. (1984). Random walks and electric networks. American Mathematical Society.

Drever, C.R., Cook-Patton, S.C., Akhter, F., Badiou, P.H., Chmura, G.L., Davidson, S.J., … Kurz, W.A. (2021). Natural climate solutions for Canada. Science Advances 7(23).

ECCC. (2013). Bird conservation strategy for Bird Conservation Region 7 in Ontario: Taiga Shield and Hudson Plains. Available from https://www.canada.ca/en/environment-climate-change/services/migratory-bird-conservation/regions-strategies/description-region-7/ontario.html.

ECCC. (2020). Canada joins the High Ambition Coalition for Nature and People. Available from https://www.canada.ca/en/environment-climate-change/news/2020/09/canada-joins-the-high-ambition-coalition-for-nature-and-people.html

ECCC. (2021). Canadian Protected and Conserved Areas Database. Available from https://www.canada.ca/en/environment-climate-change/services/national-wildlife-areas/protected-conserved-areas-database.html

Goldstein, A., Turner, W.A., Spawn, S.A., Anderson-Teixeira, K.J., Cook-Patton, S., Fargione, J., … Hole, D.G. (2020). Protecting irrecoverable carbon in Earth’s ecosystems. Nature Climate Change 10(4):287–295.

Government of Canada. (2022). 2030 Emissions Reduction Plan: Clean Air, Strong Economy Available from https://www.canada.ca/en/services/environment/weather/climatechange/climate-plan/climate-plan-overview/emissions-reduction-2030.html.

Grantham, H.S., Duncan, A., Evans, T.D., Jones, K.R., Beyer, H.L., Schuster, R., … Watson, J.E.M. (2020). Anthropogenic modification of forests means only 40% of remaining forests have high ecosystem integrity. Nature Communications 11(1).

Gunn, J.S., Ducey, M.J., & Belair, E. (2019). Evaluating degradation in a North American temperate forest. Forest Ecology and Management 432:415–426.

Hall, K.R., Anantharaman, R., Landau, V.A., Clark, M., Dickson, B.G., Jones, A., … Shah, V.B. (2021). Circuitscape in julia: Empowering dynamic approaches to connectivity assessment. Land 10(3).

Harris, L.I., Richardson, K., Bona, K.A., Davidson, S.J., Finkelstein, S.A., Garneau, M., … Ray, J.C. (2021). The essential carbon service provided by northern peatlands. Frontiers in Ecology and the Environment 20(4):222–230.

Heller, N.E., & Zavaleta, E.S. (2009). Biodiversity management in the face of climate change: A review of 22 years of recommendations. Biological Conservation 142(1):14–32.

Hirsh-Pearson, K., Johnson, C.J., Schuster, R., Wheate, R.D., & Venter, O. (2022). Canada’s human footprint reveals large intact areas juxtaposed against areas under immense anthropogenic pressure. Facets 7:398–419.

Holl, K.D., & Brancalion, P.H.S. (2020). Tree planting is not a simple solution. Science 368(6491):580–581.

Hooftman, D.A.P., Edwards, B., & Bullock, J.M. (2016). Reductions in connectivity and habitat quality drive local extinctions in a plant diversity hotspot. Ecography 39(6):583–592.

Indigenous Circle of Experts. (2018). We Rise Together: Achieving Pathway to Canada Target 1 through the creation of Indigenous Protected and Conserved Areas in the spirit and practice of reconciliation. Available from https://publications.gc.ca/collections/collection_2018/pc/R62-548-2018-eng.pdf.

IPBES. (2019). Global assessment report on biodiversity and ecosystem services of the Intergovernmental Science-Policy Platform on Biodiversity and Ecosystem Services.

IPCC. (2022a). Climate Change 2022: Impacts, Adaptation, and Vulnerability (Contribution of Working Group II to the Sixth Assessment Report of IPCC).

IPCC. (2022b). Climate Change 2022: Mitigation of Climate Change (Contribution of Working Group III to the Sixth Assessment Report of the Intergovernmental Panel on Climate Change).

Kurz, W.A., Shaw, C.H., Boisvenue, C., Stinson, G., Metsaranta, J., Leckie, D., … Neilson, E.T. (2013). Carbon in Canada’s boreal forest-A synthesis. Environmental Reviews 21(4):260–292.

Lewis, S.L., Wheeler, C.E., Mitchard, E.T.A., & Koch, A. (2019). Restoring natural forests is the best way to remove atmospheric carbon. Nature 568(7750).

Locke, H., Ellis, E.C., Venter, O., Schuster, R., Ma, K., Shen, X., … Watson, J.E.M. (2019). Three global conditions for biodiversity conservation and sustainable use: An implementation framework. National Science Review 6(6):1080–1082.

Malcolm, J.R., Holtsmark, B., & Piascik, P.W. (2020). Forest harvesting and the carbon debt in boreal east-central Canada. Climatic Change 161(3):433–449.

Marchese, C. (2015). Biodiversity hotspots: A shortcut for a more complicated concept. Global Ecology and Conservation 3:297–309.

Marrotte, R.R., Bowman, J., Brown, M.G.C., Cordes, C., Morris, K.Y., Prentice, M.B., & Wilson, P.J. (2017). Multi-species genetic connectivity in a terrestrial habitat network. Movement Ecology 5(1):1–11.

Maxwell, S.L., Cazalis, V., Dudley, N., Hoffman, M., Rodrigues, A.S.L., Stolton, S., … Watson, J.E.M. (2020). Area-based conservation in the twenty-first century. Nature 586(7828):217–227.

McRae, B.H., & Beier, P. (2007). Circuit theory predicts gene flow in plant and animal populations. Proceedings of the National Academy of Sciences of the United States of America 104(50):19885–19890.

Mitchell, M.G.E., Schuster, R., Jacob, A.L., Hanna, D.E.L., Dallaire, C.O., Raudsepp-Hearne. C., … Chan, K.M.A. (2021). Identifying key ecosystem service providing areas to inform national-scale conservation planning. Environmental Research Letters 16(1).

Morelli, T.L., Barrows, C.W., Ramirez, A.R., Cartwright, J.M., Ackerly, D.D., Eaves, T.D., … Thorne, J.H. (2020). Climate-change refugia: biodiversity in the slow lane. Frontiers in Ecology and the Environment 18(5):228–234.

Murray, D.L., Peers, M.J.L., Majchrzak, Y.N., Wehtje, M., Ferreira, C., Pickles, R.S.A., … Thornton, D.H. (2017). Continental divide: Predicting climatemediated fragmentation and biodiversity loss in the boreal forest. PLoS ONE 12(5).

NCC. (2021). Alfred Bog. Available from https://www.natureconservancy.ca/en/where-we-work/ontario/featured-projects/alfred-bog.html.

Nesshöver, C., Assmuth, T., Irvine, K.N., Rusch, G.M., Waylen, K.A., Delbaere, B., … Wittmer, H. (2017). The science, policy and practice of nature-based solutions: An interdisciplinary perspective. Science of the Total Environment 579:1215–1227.

Noon, M.L., Goldstein, A., Ledezma, J.C., Roehrdanz, P.R., Cook-Patton, S.C., Spawn-Lee, S.A., … Turner, W.A. (2021). Mapping the irrecoverable carbon in Earth’s ecosystems. Nature Sustainability 5(1):37–46.

Noss, R.F., Dobson, A.P., Baldwin, R., Beier, P., Davis, C.R., Dellasala, D.A., … Tabor, G. (2012). Bolder Thinking for Conservation. Conservation Biology 26(1):1–4.

NRCAN. (2015). Alien, invasive hemlock woolly adelgid found in Ontario. Available from https://publications.gc.ca/collections/collection_2015/rncan-nrcan/Fo123-1-114-eng.pdf.

OMECP. (2020). Woodland Caribou Recovery Strategy. Available from https://www.ontario.ca/page/woodland-caribou-recovery-strategy.

OMNRF. (2019). Woodland Caribou and the Far North. Available from https://www.ontario.ca/page/woodland-caribou-and-far-north.

OMNRF. (2021). State of Ontario’s Natural Resources – Forestry 2021. Available from https://www.ontario.ca/page/state-ontarios-natural-resources-forest-2021#:~:text=I am pleased to report,today and for future generations.

Ontario Parks. (2021). State of Ontario’s Protected Areas Indicator Report. Available from https://www.ontarioparks.com/pdf/sopar/SOPAR_ProtectedAreasSummary.pdf.

Packalen, M.S., Finkelstein, S.A., & McLaughlin, J.W. (2014). Carbon storage and potential methane production in the Hudson Bay Lowlands since mid-Holocene peat initiation. Nature Communications 5(4078).

Phillips, P., Clark, M.M., Baral, S., Koen, E.L., & Bowman, J. (2021). Comparison of methods for estimating omnidirectional landscape connectivity. Landscape Ecology 36(6):1647–1661.

Pither, R., O’Brien, P., Brennan, A., Hirsh-Pearson, K., & Bowman, J. (2021). Areas Important for Ecological Connectivity Throughout Canada. bioRxiv DOI: https://doi.org/10.1101/2021.12.14.472649.

Poley, L.G., Schuster, R., Smith, W., & Ray, J.C. (2022). Identifying differences in roadless areas in Canada based on global, national, and regional road datasets. Conservation Science and Practice 4(4):e12656.

Price, D.T., Alfaro, R.I., Brown, K.J., Flannigan, M.D., Fleming, R.A., Hogg, E.H., … Venier, L.A. (2013). Anticipating the consequences of climate change for Canada’s boreal forest ecosystems. Environmental Reviews 21(4).

Proctor, C.A., Schuster, R., Buxton, R.T., & Bennett, J. (2022). Prioritization of public and private land to protect species at risk habitat. Conservation Science and Practice. https://doi.org/10.1111/csp2.12771.

Quinby, P. (2020). The Catchacoma Ancient Forest Landscape: An Initial Inventory of Species and Habitats. Available from http://www.ancientforest.org/wp-content/uploads/ResRept39-SppHabitatInventory_PQ_APR10-2020_final.pdf.

Schloss, C.A., Cameron, D.R., McRae, B.H., Theobald, D.M., & Jones, A. (2022). “No-regrets” pathways for navigating climate change: planning for connectivity with land use, topography, and climate. Ecological Applications 32(1):e02468.

Schuster, R., Germain, R.R., Bennett, J.R., Reo, N.J., & Arcese, P. (2019). Vertebrate biodiversity on indigenous-managed lands in Australia, Brazil, and Canada equals that in protected areas. Environmental Science and Policy 101:1–6.

Secretariat of the CBD. (2020). Update of the Zero Draft of the Post-2020 Global Biodiversity Framework. Available from https://www.cbd.int/doc/c/3064/749a/0f65ac7f9def86707f4eaefa/post2020-prep-02-01-en.pdf.

Seddon, N., Chausson, A., Berry, P., Girardin, C.A.J., Smith, A., & Turner, B. (2020). Understanding the value and limits of nature-based solutions to climate change and other global challenges. Philosophical Transactions of the Royal Society B: Biological Sciences 375(1794).

Seddon, N., Smith, A., Smith, P., Key, I., Chausson, A., Girardin, C., … Turner, B. (2021). Getting the message right on nature-based solutions to climate change. Global Change Biology 27(8):1518–1546.

Smith, R.J., Bennun, L., Brooks, T.M., Butchart, S.H.M., Cuttelod, A., Di Marco, M.,… Scaramuzza, C.A. de M. (2019). Synergies between the key biodiversity area and systematic conservation planning approaches. Conservation Letters 12(1):e12625.

Smith, W., & Cheng, R. (2016). Canada’s intact forest landscapes updated to 2013. Ottawa.

Sothe, C., Gonsamo, A., Arabian, J., Kurz, W.A., Finkelstein, S.A., & Snider, J. (2022). Large Soil Carbon Storage in Terrestrial Ecosystems of Canada. Global Biogeochemical Cycles 36(2):e2021GB007213.

Soto-Navarro, C., Ravilious, C., Arnell, A., De Lamo, X., Harfoot, M., Hill, S.L.L., … Kapos, V. (2020). Mapping co-benefits for carbon storage and biodiversity to inform conservation policy and action. Philosophical Transactions of the Royal Society B: Biological Sciences 375(1794).

Stralberg, D., Arseneault, D., Balzter, J.L., Barber, Q.E., Bayne, E.M., Boulanger, Y., … Whitman, E. (2020). Climate-change refugia in boreal North America: what, where, and for how long? Frontiers in Ecology and the Environment 18(5):261–270.

Stralberg, D., Carroll, C., & Nielsen, S.E. (2020). Toward a climate-informed North American protected areas network: Incorporating climate-change refugia and corridors in conservation planning. Conservation Letters 13(4).

Thompson, P.L., Rayfield, B., & Gonzalez, A. (2017). Loss of habitat and connectivity erodes species diversity, ecosystem functioning, and stability in metacommunity networks. Ecography 40(1):98–108.

Townsend, J., Moola, F., & Craig, M.K. (2020). Indigenous peoples are critical to the success of nature-based solutions to climate change. Facets 5(1):551–556.

USEPA. (2015). Greenhouse Gas Equivalencies Calculator. Available from https://www.epa.gov/energy/greenhouse-gas-equivalencies-calculator.

Valach, A.C., Kasak, K., Hemes, K.S., Anthony, T.L., Dronova, I., Taddeo, S., … Baldocchi, D.D. (2021). Productive wetlands restored for carbon sequestration quickly become net CO2sinks with site-level factors driving uptake variability. PLoS ONE 16(3).

Wang, C., Zhang, W., Li, X., & Wu, J. (2021). A global meta-analysis of the impacts of tree plantations on biodiversity. Global Ecology and Biogeography 31(3):576–587.

Ward, M., Saura, S., Williams, B., Ramírez-Delgado, J.P., Arafeh-Dalmau, N., Allan, J.R., … Watson, J.E.M. (2020). Just ten percent of the global terrestrial protected area network is structurally connected via intact land. Nature Communications 11(1).

Watson, J.E.M., Evans, T., Venter, O., Williams, B., Tulloch, A., Stewart, C., … Lindenmayer, D. (2018). The exceptional value of intact forest ecosystems. Nature Ecology and Evolution 2(4):599–610.

Wells, J.V., Dawson, N., Culver, N., Reid, F.A., & Morgan Siegers, S. (2020). The State of Conservation in North America’s Boreal Forest: Issues and Opportunities. Frontiers in Forests and Global Change 3.

Zhu, L., Hughes, A.C., Zhao, X.Q., Zhou, L.J., Ma, K.P., Shen, X.L., … Watson, J.E.M. (2021). Regional scalable priorities for national biodiversity and carbon conservation planning in Asia. Science Advances 7(35).

