## Supplemental Figure 1 for "Integrating carbon stocks and wildlife connectivity for nature-based climate solutions"

### Supplementary Material

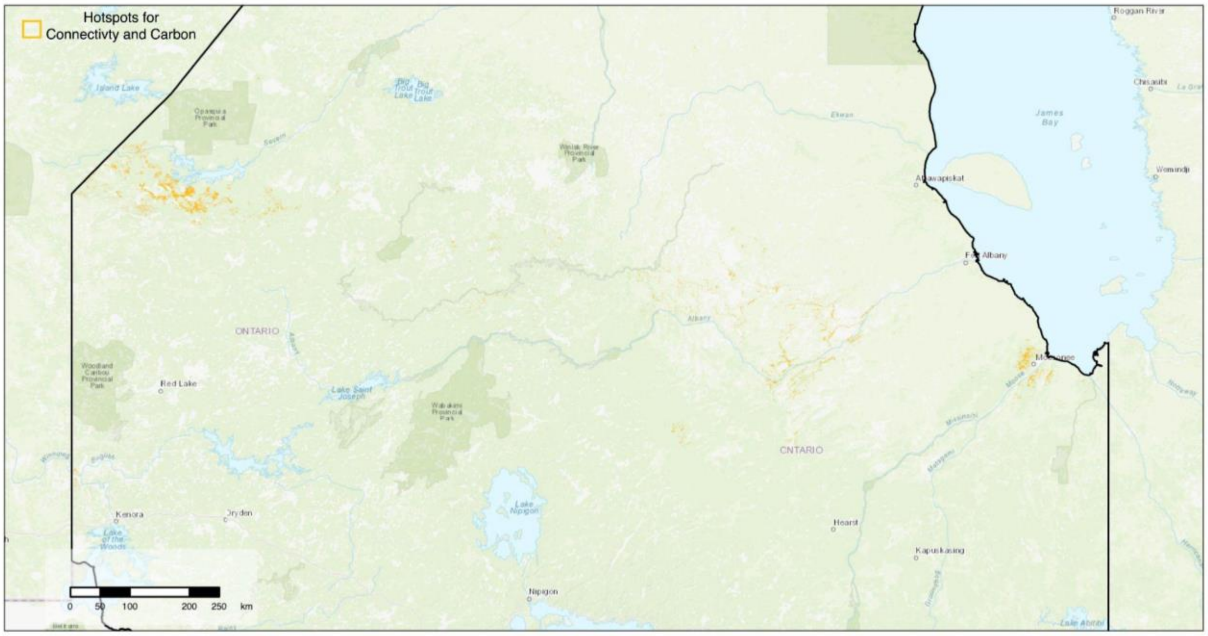


**Fig. S1.** Hotspots for terrestrial connectivity, forest carbon, and soil carbon in Ontario, Canada. Yellow regions represent 300x300m raster cells that are in the top 20% quantile for current density, forest carbon and soil carbon.
